# Synphilin-1 as a modulator of aSyn assembly

**DOI:** 10.1101/2024.03.05.583522

**Authors:** Diana F. Lázaro, Triana Amen, Ellen Gerhardt, Patrícia I. Santos, Dragomir Milovanovic, Günter Höglinger, Tiago F. Outeiro

**Affiliations:** Department of Pathology and Laboratory Medicine, Institute on Aging and Center for Neurodegenerative Disease Research, University of Pennsylvania School of Medicine, Philadelphia, PA, USA; Global Health Institute, École Polytechnique Fédérale de Lausanne, Lausanne, Switzerland; School of Biological Sciences, University of Southampton, Southampton, UK; Department of Experimental Neurodegeneration, Center for Biostructural Imaging of Neurodegeneration, University Medical Center Göttingen, Göttingen, Germany; Laboratory of Molecular Neuroscience, German Center for Neurodegenerative Diseases (DZNE), Berlin, Germany; Einstein Center for Neuroscience, Charité-Universitätsmedizin Berlin, Corporate Member of Freie Universität Berlin, Humboldt-Universität Berlin, and Berlin Institute of Health, 10117 Berlin, Germany; Department of Neurology, University Hospital, Ludwig-Maximilians-Universität (LMU), Munich, Germany; German Center for Neurodegenerative Diseases (DZNE), Munich, Germany; Munich Cluster for Systems Neurology (SyNergy), Munich, Germany; Max Planck Institute for Multidisciplinary Science, Göttingen, Germany; Translational and Clinical Research Institute, Faculty of Medical Sciences, Newcastle University, Framlington Place, Newcastle Upon Tyne, NE2 4HH, UK; Scientific employee with an honorary contract at Deutsches Zentrum für Neurodegenerative Erkrankungen (DZNE), Göttingen, Germany

**Author notes:** **Corresponding authors** Dr. Diana F. Lázaro Center for Neurodegenerative Disease Research, Department of Pathology and Laboratory Medicine, Perelman School of medicine at the University of Pennsylvania 3600 Spruce St, 3rd Floor Maloney Building Philadelphia, PA 19104, USA, Prof. Dr. Tiago F. Outeiro Department of Experimental Neurodegeneration, University Medical Center Göttingen, 37073 Göttingen, Germany.

**Keywords:** Synphilin-1, alpha-synuclein, aggregation, Parkinson’s disease, proteostasis

## Abstract

Alpha-synuclein (aSyn) is an intrinsically disordered protein that undergoes phase-separation and is associated with several neurodegenerative conditions. However, the function and the pathological role of aSyn are still elusive. Here, we modeled different types of aSyn assemblies in living cells, and developed a model that reports on gel and solid-like inclusions based on the coexpression of aSyn and synphilin-1 (Sph1). We identified striking morphological differences between aSyn-aSyn and Sph1-aSyn assemblies, characterized by distinct antibody recognition patterns, resistance to Proteinase K treatment, and protein mobilities. Importantly, we showed that the interaction between Sph1-aSyn can be manipulated, altering inclusion size and number. Sph1-aSyn interactions were central for inclusion formation and localization, and that inclusions include lysosomes and AP-1 vesicles, consistent with previous studies in human brain tissue. In total, we provide novel insight into the biology of protein aggregation, shedding light on potential therapeutic strategies that extend beyond conventional targets. Deciphering the role of Sph1 and other aSyn-interacting proteins on aSyn biology and pathobiology will be essential for treating synucleinopathies.

## Introduction

Synphilin-1 (Sph1) was first identified as an alpha-synuclein-interacting (aSyn) protein in yeast two-hybrid screening ^1^. The interaction was validated by fluorescence resonance energy transfer in cultured cells ^2^, and co-expressing of the Non-amyloid component (NAC) domain of aSyn with Sph1 resulted in the formation of cytoplasmic inclusions ^1, 3^. Sph1 is expressed in various brain regions, including the *substantia nigra*, hippocampal pyramidal, cerebellar Purkinje cells and glial cells ^4, 5, 6^, and is also enriched in the heart and placenta ^1^. The function of Sph1 is still unclear, but several studies connect it to degradation pathways ^3^, as well as to synaptic vesicle-binding protein ^5, 6^. The accumulation of Sph1 in Lewy bodies (LBs) and in glial cytoplasmatic inclusions (GCIs) has been reported ^7, 8^, but its contribution to the formation of these structures, remains unclear. Notably, Sph1 is found within the central core of the LBs while aSyn is more peripheral ^6, 7, 9^. The Sph1 sequence exhibits several conserved domains including ankyrin-like repeats and a coiled-coil domain, known to be involved in protein-protein interactions, ^4^. Therefore, it is not surprising that Sph1 interacts with several proteins such as Parkin, SIAH, LRRK2, and PINK1 ^3, 10, 11, 12, 13, 14, 15, 16^. All of these proteins have been implicated in Parkinson’s disease (PD), a neurodegenerative disorder associated with the misfolding and aggregation of aSyn.

aSyn, is considered to play a major role in the etiology of PD ^17, 18, 19, 20^. This intrinsically disordered protein predominantly localizes at the presynaptic terminals and nucleus ^21, 22, 23^. Functionally, aSyn is involved in SNARE-complex assembly, and in synaptic function. However, whether and how aSyn acquires a toxic function remains poorly understood ^24, 25, 26^. Most likely the process involves a combination of factors, such as the decline of cellular proteostasis and/or intrinsic modifications within the protein (such as post-translational modifications) ^27, 28, 29, 30^. The gain of toxicity by aSyn assemblies is also thought to play a role in pathogenesis. Yet, studying the early steps of aggregation has been challenging. For instance, high concentrations of monomeric aSyn are required for the nucleation process to occur *in vitro*, and in living cells, seeding or chemical treatments (such as rotenone, MPTP, protein degradation blockers) are typically used to elicit aSyn aggregation ^31, 32, 33, 34, 35, 36^. Intriguingly, aSyn does not easily form inclusions upon overexpression in cells or in animal models.

Here, we developed a new simple cell-based model to investigate the role of Sph1 on aSyn biology/pathobiology. Our model allowed us to monitor the formation of aSyn-Sph1 assemblies in living cells, as a result of a biologically-relevant protein-protein interaction. We demonstrated that aSyn-aSyn and Sph1-aSyn assemblies are morphologically different to each other due to (i) different antibody recognition under native condition, (ii) distinctive resistances of aSyn to Proteinase K (PK) treatment, and (iii) different protein mobility in protein electrophoresis. Furthermore, we show that the protein interactions are important for localization and inclusion formation. By manipulating the interacting region, or expressing different protein ratios, we altered the propensity and the phenotype of Sph1-aSyn inclusions. Overall, our results indicate that Sph1 is a key factor in the regulation of aSyn protein aggregation.

## Results

### Sph1-aSyn interaction promotes the formation of cytoplasmic inclusions

To investigate the role of Sph1 on aSyn aggregation, we generated a new Bimolecular Fluorescence Complementation (BiFC) assay for Sph1: VN-Sph1 and Sph1-VC (henceforth referred to as Sph1 BiFC; more information in Materials and Methods section). The two halves of Venus fluorescence protein can be reconstituted upon protein-protein interaction, allowing the direct visualization of dimers/oligomers in living cells. As we previously reported, in HEK 293 (HEK) cells co-transfected with VN-aSyn and aSyn-VC (henceforth referred to as aSyn BiFC) typically shows a diffuse/homogeneous distribution of Venus signal throughout the cytoplasm and nucleus ^37^ (with some exceptions ^38^). This has been a limitation in the field, as aSyn expression in cell models typically does not form visible aggregates (Figure 1A, top row). In contrast to aSyn BiFC, Sph1 BiFC formed inclusions with round and elongate/tubular morphology (Figure 1B, top row). Immunostaining the cells against Sph1, revealed not only the inclusions but also some soluble Sph1 within the cells, indicating that the formation of Sph1 inclusions is correlated with protein-protein interaction and with the orientation of the Sph1 molecules (Figure 1B, top row). Importantly, Sph1 expression also resulted in the formation of cytoplasmic inclusions (Figure 1B, lower panel-Sph1-V5), demonstrating that Sph1 aggregates on its own, independently of the Venus fragment. Furthermore, expressing VN-Sph1 + VC or VN + Sph1-VC did not produce fluorescent signal (supplementary data Figure 1A). Next, we examined the effects of co-expressing Sph1 with aSyn. The interaction occurred, but depended on the orientation of the proteins. The signal was detected in cells co-expressing VN-Sph1 + aSyn-VC after 24 hours (Figure 1A, supplementary data Figure 1B), but only a weak signal was detected in cells co-expressing VN-aSyn + Sph1-VC after 48 hours (Figure 1A). Moreover, only VN-Sph1 + aSyn-VC formed visible cytoplasmic inclusions (Figure 1A and 1B, middle panel). Then, we asked whether phosphorylation would play a role on inclusion formation. To test this, we used mutants that block (S129A) or mimic (S129D) phosphorylation. No differences were observed in terms of Sph1-aSyn inclusions (supplementary data Figure 1C), suggesting that phosphorylation does not interfere with Sph1 and aSyn interaction, nor with the formation of this type of cytosolic inclusions. We also challenged the system by expressing first aSyn BiFC and, after 24 hours, introducing VN-Sph1 or Sph1-VN (Figure 1D). These experiments did not result in the formation of visible inclusions, suggesting that the initial interaction is a key factor for aSyn assembly. We also applied this new inclusion-formation model in primary cortical and hippocampal neurons (Figure 1D and supplementary Figure 1C). As observed in HEK cells, Sph1-aSyn formed cytosolic inclusions in both cortical and hippocampal neurons. This suggests that the formation of Sph1-aSyn inclusions occurs in different cell types, including primary neuronal cells. However, due to limitations in transfection rates and/or protein expression, we used the HEK cell line for the remainder of our study. Next, we assessed the protein levels using Western blot (WB) analyses (Figure 1E). To account for the molecular weight differences between aSyn and Sph1, we ran separate gels for Sph1 BiFC to resolve VN-Sph1 and Sph1-VC (Figure 1E, right panel). Overall, we observed lower levels of aSyn-VC, which limited some of the assays performed. However, it is important to note that these observations are not indicative of the degradation of aSyn-VC, as our group and others have reported similar observation ^38, 39, 40, 41, 42, 43, 44^. Although it may appear unexpected, we view this distinction as biologically significant, given that protein levels in cells are affected by a multitude of factors, thereby contributing to biological diversity.

**Figure 1.**
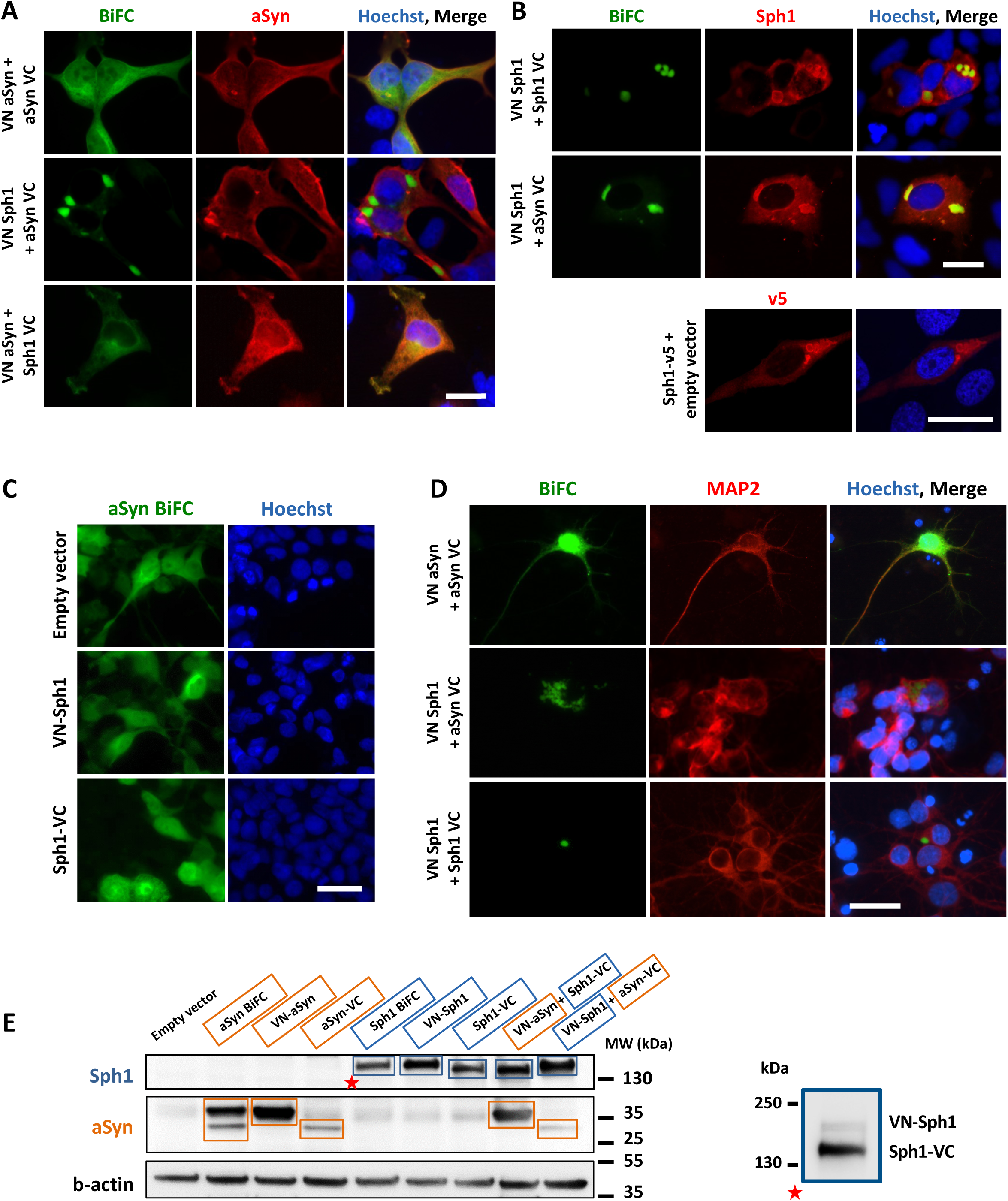
Sph1-aSyn form cytoplasmic inclusions in HEK 293 cells and in primary cortical neurons. **A-B. Immunocytochemistry of Sph1-aSyn inclusions.** HEK 293 cells were fixed and stained using either anti-Sph1 or anti-V5 antibodies (A) or total aSyn (Syn1) (B) (scale bar: 30 µm). **C. Competition assay.** HEK293 cells were initially transfected with aSyn BiFC, and after 24 hours, the cells were again transfected with empty vector, VN-Sph1 or Sph1. No inclusions were observed, indicating that Sph1 is important to initiate the aggregation of aSyn. **D. Primary cortical neurons exhibit Sph1-aSyn inclusions.** Transfected primary cortical neurons at DIV4, exhibit similar inclusions after 48 hours (DIV6) showing that the model works independently of the cell type (scale bar: 30 µm). **E. Expression levels of aSyn and Sph1.** Immunoblot analyses of the levels of aSyn (orange box) and Sph1 (blue box). A separate blue square shows a longer run for Sph1 BiFC, displaying the separation of both VN-Sph1 and Sph1-VC. All the above experiments were performed at least three times.

To the best of our knowledge, this is the first time that the interaction of two proteins found in the LBs, Sph1 and aSyn, could be observed in living cells without the need for fixation and immunostaining. Overall, these results suggest that the initial interaction plays a crucial role in aSyn assembly and requires a specific orientation of the proteins to occur.

### Sph1-aSyn inclusions do not impact cellular homeostasis

We next evaluated the toxicity of Sph1-aSyn using different readouts. We measured the release of adenylate kinase as an indicator of membrane integrity (Figure 2A), and assessed cell viability using two different assays: cell proliferation (Figure 2B) and cell viability (Figure 2C). Overall, none of these readouts showed signs of toxicity and/or viability impartments, suggesting that the accumulation of Sph1-aSyn is not, *per se*, detrimental for cellular homeostasis. Yet, it has been widely reported that aSyn has detrimental implications for different subcellular compartments, such as the Golgi apparatus in the *substantia nigra* ^45^. Normally, when the Golgi apparatus is labeled with antibodies like anti-giantin, it appears as a perinuclear formation, either compact or elongated. However, under stress conditions, the Golgi complex exhibits an increase in discrete Golgi objects and eventually breaks into smaller fragments distributed throughout the cytoplasm ^46, 4748^. Therefore, we assessed the number of transfected and non-transfected cells with normal, diffuse, and fragmented Golgi (Figure 2D and E, supplementary data Figure 2A). Similar to our previous observation both in HEK cells and in neuronal Lund Human Mesencephalic (LUHMES) cells ^40,49^, expression of aSyn BiFC constructs increased Golgi fragmentation (Figure 2D and E and supplementary Figure 2A, red bars) in comparison to the control. These results can be associated to aSyn expression since non-transfected cells did not exhibit the same phenotype. Nevertheless, Sph1 counter-balanced the effects of aSyn, as the expression of both proteins reduced Golgi fragmentation by ≍20% in comparison to aSyn-aSyn (Figure 2D). In order to further validate our observations, we synchronized the cultures by inducing cell cycle arrest at the G1/S boundary by implementing a double-thymidine S phase block ^50^. After the removal of the S phase block through thymidine washout, and expressing aSyn and Sph1 (additional details in the Materials and Methods section) HEK cells were fixed 12 hours after the thymidine removal, and assess the Golgi apparatus morphology as the before. Overall, we did not observe any differences between untreated cells (not synchronized) and cells treated with thymidine (at the G1/S phase), indicating that the effects previously observed were indeed due to αSyn expression rather than the cell cycle stage (supplementary Figure 2B). We then analyzed the area occupied by the Golgi apparatus. The rationale behind this was that fragmented Golgi would occupy a larger area as the smaller fragments are dispersed throughout the cytoplasm, thus covering a greater surface area, compared to the normal and diffuse Golgi. Due to the irregular and diverse shapes that Golgi can assume, we used a plugin from ImageJ (Hull and Circle) to extract some shape descriptors as the convex area (supplementary Figure 2C). We selected the convex area data as the best approximated to the area occupied by the Golgi. As illustrated by the frequency distribution, we observed that the percentage of Golgi area for cells expressing aSyn BiFC is lower for smaller areas, but increases with the larger areas (supplementary Figure 2C, orange bars). In contrast, VN-Sph1 + aSyn-VC displayed a frequency distribution more similar to that observed with the control cells. This semi-automatic quantification aligned with our quantifications, indicating that aSyn BiFC, occupies a larger area compared with control or VN-Sph1 + aSyn-VC. Overall, these results suggest that the interaction between both proteins is beneficial, reducing proteostasis stress.

**Figure 2.**
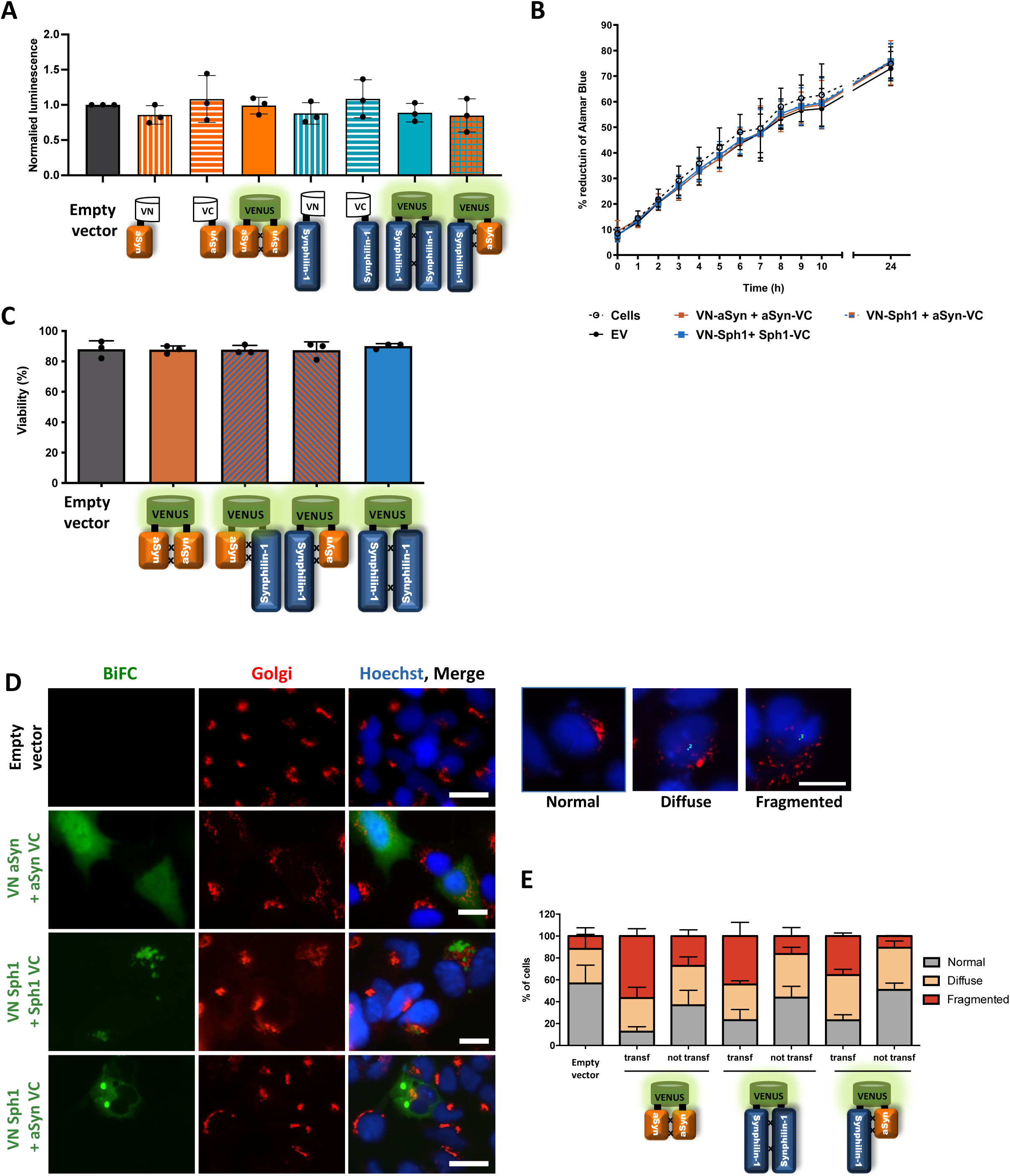
Sph1 rescues the detrimental effect of aSyn on the Golgi apparatus. **A-C. Sph1-aSyn inclusions do not induce toxicity. A. Measurement of adenylate kinase (AK) release to the cell medium.** The medium collected from the cells was used to measure AK activity as a measure of cytotoxicity for each condition. **B. Quantification of the metabolic activity of the cells.** We also investigated if any of the tested conditions had an impact on the proliferation of the cells, but no major differences were observed. **C. Evaluation of cell membrane integrity.** Cell death was evaluated by measuring the viability of the cells (trypan blue quantification). None of the conditions showed differences in terms of viability or cells number. All measurements were performed 24 hours after expression of Sph1 and aSyn **D-E. Sph1 rescues the Golgi fragmentation phenotype induced by aSyn.** 48 hours after transfection, the morphology of the Golgi apparatus was analyzed and quantified. We observed that aSyn-aSyn displayed an increased Golgi fragmentation compared to Sph1-Sph1 and Sph1-aSyn. Importantly, Sph1 rescues this phenotype by decreasing the amount of fragmented Golgi by 40%. Also, these results can be attributed to aSyn expression since the surrounding cells did not present the same fragmentation degree (scale bar: 30 µm). All the experiments were performed in triplicates.

### Characterization of Sph1-aSyn assemblies

To understand the biogenesis of Sph1-aSyn inclusions, we used several biochemical assays to measure the protein aggregation state and stability. We started by analyzing the properties of Sph1-aSyn complexes in native conditions, by using different aSyn antibodies. For Sph1-aSyn, we did not detect any band using the MJF 14-6-4-2 antibody (conformation-specific antibody that detects aggregated aSyn), and only faint bands for Syn1 (that detects total aSyn) and 5G4 antibody (recognizing aggregated aSyn) (an upper band located at the well region) (Figure 3A). As stated earlier, aSyn-VC accumulates at lower levels when compared to VN-aSyn (Figure 1C). Therefore, either the amount of aSyn present in the sample is below the detection range of the assay or aSyn is not fibrillized. To further characterize the nature of these inclusions, we performed size exclusion chromatography by high-performance liquid chromatography (SEC-HPLC) followed by dot blot analysis, and we detected aSyn using the Syn1 antibody. Co-expression of the proteins, either VN-aSyn + Sph1-VC or VN-Sph1 + aSyn-VC, did not significantly alter the high molecular weight of aSyn species compared with the controls (VN-or -VC) (Figure 3B). Instead, Sph1-aSyn formed small assemblies (Figure 3B).

**Figure 3.**
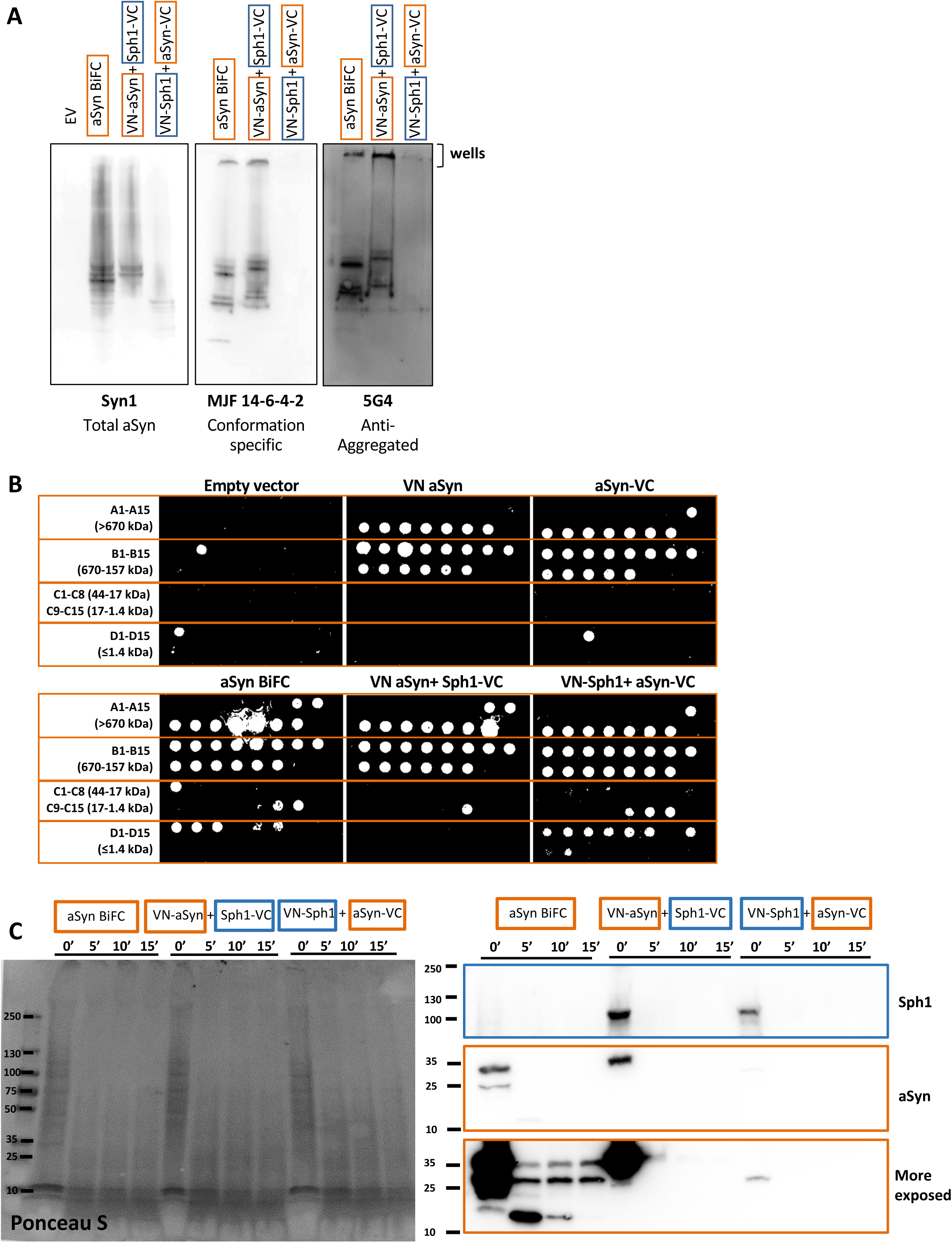
Biochemical characterization of Sph1-aSyn inclusions. **A-B. At the molecular level, Sph1-aSyn induces slight changes on aSyn.** Under non-denaturing condition, a smear for total aSyn was visible for all conditions, indicating the presence of oligomeric species. The used of different antibodies show lower, or no signal suggesting that either Sph1-aSyn does not form high molecular species or, due to the low expression level of aSyn-VC, it is not possible to be detected such species. Nevertheless, a band at the wells regions was detected for 5G4, an antibody that recognizes aggregated aSyn. n = 2 (A). HPLC/dot blot assay did not show major shifts in molecular weight of aSyn. These data suggest that the inclusions observed might represent a phase transition or/and an amorphous state. n = 2. **C. Sph1-aSyn inclusion are less resistant to PK than aSyn-aSyn.** Protein lysates were digested with PK for different times (0, 5, and 10 min). Overall, Sph1-aSyn and aSyn-Sph1 are less resistant to PK. n = 3.

Next, we assessed the structural stability of these assemblies by treating the samples with PK. We found that the aSyn-aSyn interaction is resistant to PK digestion, resulting in lower fragments around 15kDa. The PK resistance patterns for both aSyn-Sph1 and Sph1-aSyn were similar, and collectively, they were less-resistant to PK digestion compared to aSyn BiFC (Figure 3C). The resistance is independent of aSyn levels, as in both cases, Sph1-aSyn and aSyn-Sph1 (where VN-aSyn construct is high expressed) the digestion was completed within 10 minutes, and no bands were detected. Collectivity, these data showed slight changes of aSyn at the molecular level, challenging our initial premise that, instead of being in the presence of aggregated clusters, Sph1-aSyn may forms amorphous inclusions.

### Sph1-aSyn inclusions display gel-like properties

Several intrinsically disordered proteins like TAR DNA-binding protein 43 (TPD 43), Fused in sarcoma (FUS) and Tau are involved in phase separation processes ^51^. Recent studies have shown that aSyn can form liquid condensates through phase separation, and that liquid–liquid phase separation of aSyn occurs before its aggregation ^52^. Therefore, we next asked whether these inclusions may represent a phase-separated liquid droplet. To investigate this, we used Fluorescence Recovery After Photobleaching (FRAP) as a useful tool to study protein state/mobility in living cells. We analyzed the recovery data from both cytoplasm and/or inclusions over the course of 5 minutes for aSyn-aSyn, Sph1-aSyn, and Sph1-Sph1 (Figure 4A and B). The fastest dynamics were observed for aSyn-aSyn (data not shown due to complete fluorescence recovery being outside of the photodetector sensitivity limit), suggesting the presence of a soluble, freely diffusible state of aSyn-aSyn species. The FRAP recovery curves for Sph1-aSyn inclusions (Figure 4B, orange circle) exhibited a less pronounce half-life recovery (supplementary data Figure 3A orange graph) when comparing to Sph1-aSyn diffuse signal areas (Figure 4B, black triangle), suggesting a different organization within the inclusions. In contrast, Sph1 BiFC assemblies remained immobile, with almost no signal recovery after the photobleaching. This indicates that the interaction between Sph1 and aSyn alters the proprieties of both proteins. These findings suggest that Sph1-aSyn can adopt a specific conformation typically described as a gel-like state.

**Figure 4.**
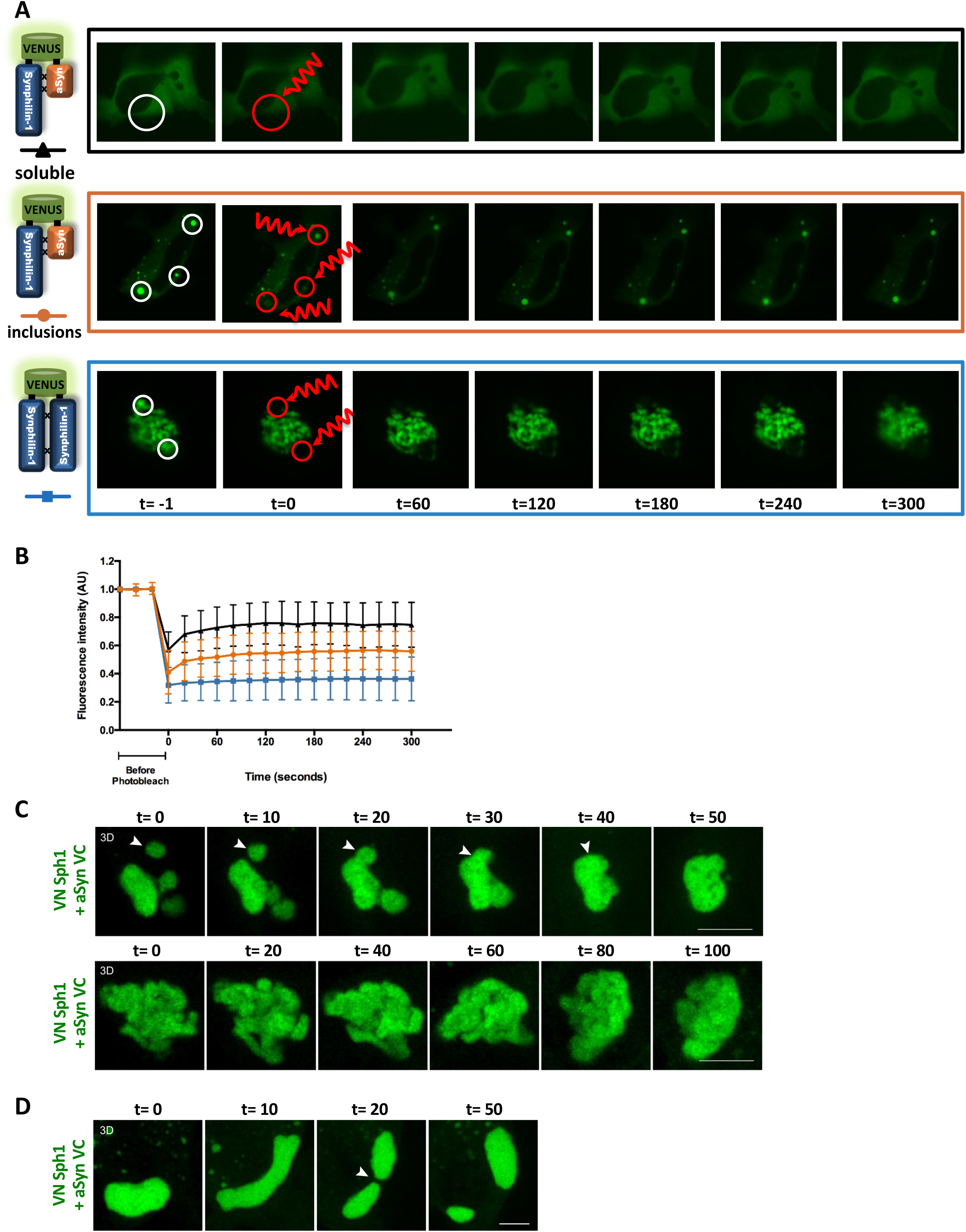
Sph1-aSyn assemblies have gel-like properties. **A-B Sph1-aSyn inclusions have a slower rate of recovery compared to aSyn-aSyn.** FRAP was performed to measure the mobility of the inclusions. Sph1-aSyn inclusions showed a slower rate of recovery compared to aSyn-aSyn inclusions, indicating that Sph1-aSyn has a more gel-like state. **C-D. Dynamics of Sph1-aSyn inclusions.** Live cell imaging was performed to determine the dynamics of Sph1-aSyn inclusions. Small or single inclusions fused to a larger aggregate, while tubular inclusions were less mobile (C). Treating the cells with cycloheximide made the inclusions more fluid, allowing for fusion and fission events to occur (D). However, after 24 hours of treatment, the inclusions were not completely dissolved (scale bar: 5 µm). **Movie 4C upper.** Illustration of the Sph1-aSyn inclusions dynamics, highlighting the fusion of small or single inclusions into larger aggregate, denoted by arrowheads. This elucidates the dynamics of Sph1-aSyn small inclusions, underscoring the processes involved in their aggregation within the cellular context. **Movie 4C lower.** The movie showcases a time-lapse sequence illustrating the decreased mobility of tubular Sph1-aSyn inclusions compared to smaller or individual inclusions. This constitutes a visually-stunning demonstration highlights the distinctive behavior of the tubular Sph1-aSyn inclusions, highlightingtheir distinct characteristics. **Movie 4D.** The cells treated with cycloheximide results in increased fluidity of the inclusions, facilitating fusion and fission events. The data highlight the dynamic nature of the inclusions and the possibility to modulate their behavior, shedding light on the underlying mechanisms involved in inclusion formation and dynamics.

Next, we performed live cell imaging to determine the dynamics of Sph1-aSyn. Generally, liquid-like condensates can grow in two different ways: by Ostwald ripening (diffusion-limited movement of molecules from small to large droplets) or by coalescence (small droplets come into close proximity, touch and then fuse into a larger droplet.)^53^. In our model, we observed two distinct morphologies: round and/or elongate/tubular inclusions. Notably, Sph1-aSyn inclusions (multiple small inclusions or single inclusions), consistently fused to form larger aggregates (Figure 4C, upper panel). The enlongated/tubular inclusions exhibited lower mobility (Figure 4C, lower panel and movie 4C upper, and lower), and did not significantly change or undergo fusion on the same time scales. Strikingly, cycloheximide treatment increased the overall fluidity of the inclusions enabling them to break apart, but it was not sufficient to completely dissolve Sph1-aSyn inclusions from the cytoplasm, even after 24 hours of treatment (Figure 4D and movie 4D). To test if these structures were modulated by oxidative stress, we induced the production of reactive oxygen species by treating the cells with sodium arsenite. Upon treatment, we observed the formation of stress granules, but no colocalization was detected, and no significant changes in Sph1-aSyn inclusions were observed (supplementary data Figure 3B). Overall, these results suggests that these Sph1-aSyn complex exist in a specific conformation resembling a gel-like state, and their mobility allow them to fuse, forming larger aggregates.

### Lysosomes and AP-1 localize within Sph1-aSyn inclusions

During our acquisitions, we frequently observed the presence of dark spots embedded within the inclusions. As several studies have shown, during aSyn aggregation, particularly in the context of pathology, these aggregates consist of a mix of organelles (such as mitochondria and lysosomes) and lipids ^54,55^. To determine whether Sph1-aSyn inclusions correlates with any cellular organelles, we monitored different membrane and non-membrane compartments during Sph1-aSyn inclusion formation (Figure 5 and supplementary data Figure 3). Interestingly, the inclusions did not co-localize with any of the examined organelles, including lipid droplets, peroxisomes, mitochondria, endoplasmic reticulum, and P bodies (supplementary data Figure 4). We observed only a slight reduction in P bodies, and some mitochondria were found to be localized proximal to the inclusions, even passing through them (Figure 5A). Otherwise, there were no significant alterations in organelle appearance, with the exception of the Golgi fragmentation, as previously described.

**Figure 5.**
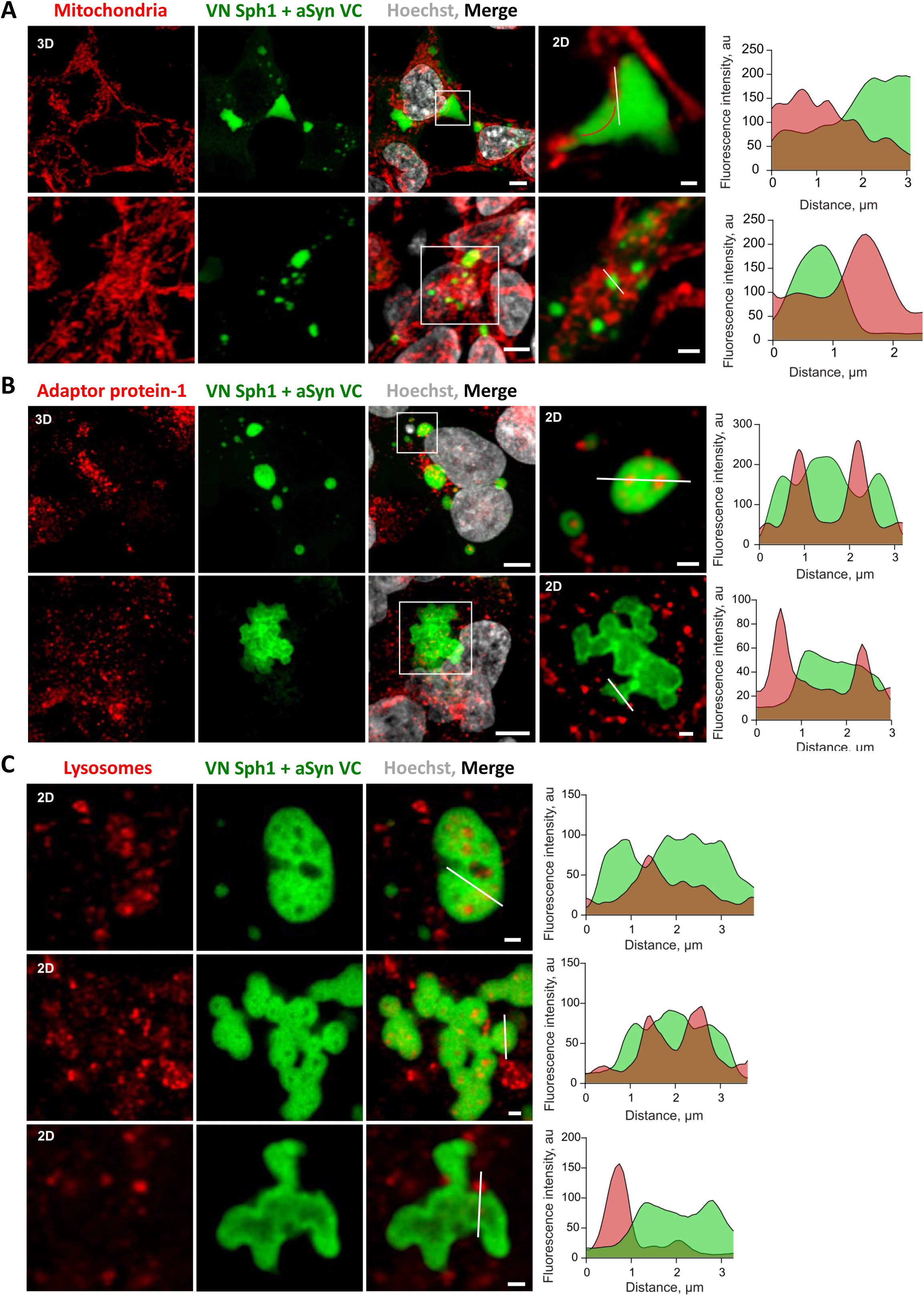
Lysosomes and AP-1 are trapped within Sph1-aSyn inclusions. We observed that mitochondria localize proximal or even passed through Sph1-aSyn (red arrow) (A), but remarkably, lysosomes and AP-1 are presented within Sph1-aSyn inclusion (B-C). Confocal imaging of HEK293T cells overexpressing Sph1-aSyn stained with TOM20 (A), AP-1 (B), or LAMP1 (C) antibodies. Representative fluorescence intensity profiles through the inclusions (white line on the inset) are shown. Scale bar of overall image: 5 µm and crop image: 1 µm.

We also noticed that inclusions were often localized proximally to the plasma membrane side of the Golgi (trans-Golgi) which serves as a membrane trafficking hub that directs vesicles towards lysosome formation or secretion. Post-Golgi compartments, but not any other compartments, were found inside the Sph1-aSyn inclusions (Figure 5B and C). Vesicles derived from the trans-Golgi (TGN) network marked with gamma-adaptin (AP-1, Adaptin γ), and lysosomes (marked with LAMP1) were found within the inclusions (but not within the tubular inclusions). This indicates that Sph1-aSyn assembly may perturbate the normal membrane trafficking originating from the TGN. These observations show that the persistent presence of these inclusions might result in compromised degradation system, such as the ubiquitin–proteasome system and/or autophagy–lysosomal pathway, as well as an impairment at the level of the TGN.

### Modulation of Sph1-aSyn inclusions by gene expression alterations and by Hsp70

As we established, the interaction between Sph1 and aSyn is important for the formation of intracellular inclusions, but the specific contribution of each protein to this process is unknown. Therefore, we asked whether Sph1 could be a driving force for our observations. To test this hypothesis, we expressed Sph1 and aSyn at different ratios. Our observations clearly showed that Sph1 expression primarily led to the formation of a single inclusion, while aSyn resulted in a more cytoplasmic distribution (Figure 6A). These results showed that Sph1 drives the assembly process and emphasizing the significance of protein-protein interactions and orientation of the proteins as essential driving force. Furthermore, independent of the ratio, we often observed Sph1-aSyn BiFC signal proximal to the plasma membrane (Figure 6B and supplementary data Figure 5A). As proof of concept, we constructed a plasmid in which we fused Sph1 and aSyn to ensure comparable expression levels of both proteins. Once again, we confirmed the presence of Sph1-aSyn at the membrane (see supplementary data Figure 4, Sph1-aSyn panel).

**Figure 6.**
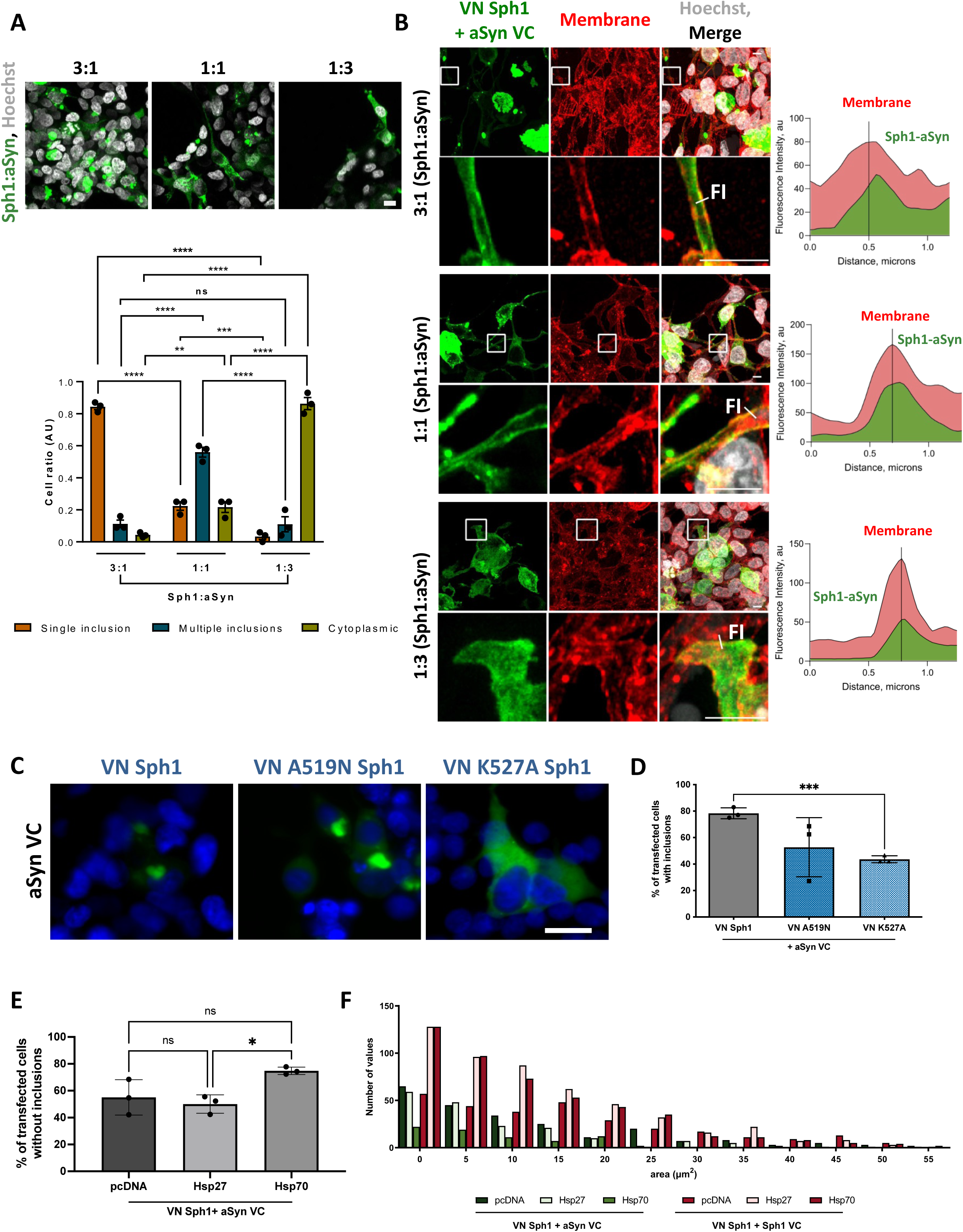
Modulation of Sph1-aSyn inclusion formation. **A. Different levels of expression of aSyn and Sph1 change the inclusion phenotype.** Higher levels of Sph1 resulted in a preferential tubular inclusion phenotype, and increasing aSyn leads to cytoplasmic localization and small multiple inclusions. **B. Sph1-aSyn localize at the membrane.** Regardless of the Sph1 and aSyn ratio, clusters of Sph1-aSyn are present within the membrane. This observation is more noticeable when stable expression of Sph1 and aSyn is promoted. Scale bar: 10 µm. **C-D. Disruption of Sph1-aSyn interaction leads to a decrease in inclusion formation.** Mutation in Sph1 region, proposed to interact with aSyn, leads to a reduction in inclusions, particularly at position 527. This shows that the interaction is important for the formation of these assemblies. Scale bar: 30 µm. **E-F. Hsp70 disaggregates Sph1-aSyn inclusions.** The expression of Hsp70 leads not only to a reduction in number but also to a reduction in size of Sph1-aSyn inclusions.

Our results suggested that Sph1 has the capacity to shift the aggregation state of aSyn. To validate the role of Sph1 on aSyn aggregation, we hypothesized that abolishing the interaction between these two proteins would result in a reduction in inclusion formation. To test this hypothesis, we strategically mutated different Sph1 residues known to be important for this interaction ^10^. We specifically targeted residues in positions 519 and 527, based on the published dissociation constants (K_D_), which reports on the strength of the interaction between a molecule and its binding partner, and because these two residues were relatively well separated from each other ^10^. The expression of these two mutants led to an overall decrease in the number of inclusions within the cells (Figure 6C and D). These results were statistically significant for K527A Sph1 mutant, with 40% of the cells displaying less inclusions in comparison with cells expressing WT Sph1 (Figure 6D). Once again, these results confirm the relevance of the Sph1 interaction in the context of aSyn assembly.

Protein aggregation is a dynamic process, and as we previously showed, overexpression of Heat shock proteins (Hsp) 70 and Hsp27 reduces aSyn toxicity and aggregation in cells and in animal models^37, 56, 57, 58^. Therefore, we sought to investigate whether Hsps would impact the interaction between Sph1 and aSyn in our new model. We observed reduction in the percentage of cells with inclusions upon Hsp70 expression (around 60% of the cells did not have inclusions) (Figure 6E). The results were also reflected in the reduction of the area size of Sph1-aSyn inclusions (Figure 6F). This reduction appears to be associated with Sph1-aSyn interaction, as we did not observe the same effect on Sph1 BiFC (Figure 6F). In fact, Hsp27 and Hsp70 led to an increase in the size of Sph1 inclusions.

Altogether, these results showed that manipulating Sph1 levels or targeting specific residues, directly interferes with aSyn assembly and localization. Our model of Sph1-aSyn aggregation is not only amenable for assessing future ways for intervening with aSyn physiology, but also reveals that Sph1-aSyn inclusion formation is a reversible and dynamic process in which Hsps, such as Hsp70, can dissolve Sph1-aSyn inclusions.

## Discussion

The mechanisms involved in the assembly of aSyn into ordered filaments that are components of intracellular inclusions are still unknown. Over the years, several systems have been developed to model the aSyn aggregation cascade in cells. However, for reasons we do not fully understand, these cellular models fail to recapitulate important aspects of aSyn aggregation. Traditionally, inhibitors, treatments with stress inducers, or external stimuli (like the addition of a recombinant protein), are used to induce aSyn aggregation. Our study demonstrates that one of the reasons why inducing aSyn aggregation may fail might be due to differences in the levels of Sph1 expression. In our new model, we showed that Sph1 contributes to aSyn aggregation based on protein-protein interactions. Importantly, we showed that we can modulate the phenotype of the inclusions formed by regulating the expression levels of both proteins or by disrupting their interaction. Not only this, protein-protein interaction represents a dynamic process that allows us to observe the fusion and fission of the inclusion, but we were also able to reduce the number and the size of Sph1-aSyn inclusions upon Hsp 70 expression. Consistent with previous studies and with our current study, the coiled-coil domain of Sph1 is important to establish the interaction with aSyn, and essential to induced inclusion formation. The data presented here, open new avenues for modulating aSyn aggregation. Unfortunately, the expression levels of Sph1 in PD patients have not been determined systematically. Hypermethylation of the *SNCAIP* gene was reported in the cortex of PD patients, which might suggest that Sph1 is down-regulated. As the authors mentioned, the sample size was small (12 cases), and all had dementia. Therefore, this suggests that hypermethylation is more likely associated with dementia in PD rather than with PD itself ^59^.

Another important observation is the localization of Sph1-aSyn at the cell membrane. We observed clusters of Sph1-aSyn at the cell membrane, but further experiments are necessary to determine whether this is caused by changes in lipid levels/chemical properties or by other cellular mechanisms triggered by Sph1 expression. Our results show that our model successfully replicates essential aspects of aSyn physiology, as the interaction between aSyn and lipids can modulate its propensity to assemble ^60^. The potential importance of membrane binding for inclusion formation and/or pathology is further supported by the presence of lysosomes and AP-1 vesicles in the inclusions. Our study demonstrates that post-Golgi compartments are sequestered into inclusions, leading to the disruption of the TGN. Interestingly, our model offers new possibilities for exploring the role of aSyn in membrane trafficking in mammalian cells, a concept previously established in the aSyn yeast model ^61^.

Overall, these results suggest that Sph1 can act as a protein-adaptor for aSyn, and possibly for other intrinsically disordered proteins. We think that Sph1, by establishing multiple interactions with proteins related with neurodegenerative diseases as PD (LRRK2, Parkin, SIAH), might play a broader and more general impact on proteostasis, perhaps as sentinel protein, which either promotes protein clearance, or compartmentalizes harmful protein. As an example from our work, not only the interaction between Sph1 and aSyn did not show signs of toxicity, but Sph1 alleviated the burden of aSyn at the Golgi apparatus. Additional studies may open novel strategies for therapeutic intervention focusing not only on the typical culprits but also on proteins such as Sph1, which we show to modulate aSyn aggregation and overall cellular proteostasis.

## Methods

### Sph1 BiFC

The cDNA sequence of human Sph1 was subcloned from the pcDNA3.1/V5-His-TOPO expression vector (Invitrogen, USA) ^62^ into the Venus-BiFC plasmids previously described ^37^. Briefly, the Sph1 sequence was cloned using specific primers 3’ of the N-terminal fragment of Venus (VN; VN-Sph1), corresponding to amino acids 1–158 (VN-fragment), and upstream of the C-terminal fragment (VC; Sph1-VC), corresponding to amino acids 159–239, using specific primers. The primers contained the restriction enzyme sites AflII at the 5’ and XhoI at the 3’-end, respectively. The sequences of the primers used were as follows:

Forward VN-Sph1: 5’-CCCCTTAAGATGGAAGCCCCTGAATACCTTG-3′

Reverse VN-Sph1: 5’-CCCCTCGAGTTATGCTGCCTTATTCTTTCCTTTGCTAGCGGAGCTGG-3′

Forward Sph1-VC: 5’-CCCCTTAAGATGGAAGCCCCTGAATACCTTG-3′

Reverse Sph1-VC: 5′-CCCCTCGAGTGCTGCCTTATTCTTTCCTTTGCTAGCGGAGCTGG-3′

The PCR fragments were digested and cloned into the aSyn BiFC constructs by replacing the aSyn insert. All constructs were verified by DNA sequencing.

To create the aSyn and Sph1 fusion, Sph1 was cloned into pcDNA3.1-aSyn-myc plasmid between Xho1 and Not1 restriction enzyme sites. aSyn-myc is inserted with Nhe1 and Xho1 restriction sites. The primers used were the follow:

Forward Sph1-xho: 5’-GATCCTCGAGGAAGCCCCTGAATACCTTGATT-3’

Reverse Sph1-not: 5’-GATCGCGGCCGCTTATGCTGCCTTATTCTTTCCTTTGC-3’

### Site directed mutagenesis on Sph1

For mutagenesis, the primers were designed according to the manufacturer’s instructions. We used the following primers:

Forward Sph1_A519N: 5’-GACCTGCATGTCGCTGAACTCTCAAGTGGTGAAG-3’

Reverse Sph1_A519N: 5’-CTTCACCACTTGAGAGTTCAGCGACATGCAGGTC-3’

Forward Sph1_K527A: 5’-GTGGTGAAGTTAACCGCGCAGCTAAAGGAAC-3’

Reverse Sph1_K527A: 5’-GTTCCTTTAGCTGCGCGGTTAACTTCACCAC-3’

Site-directed mutagenesis was performed using the QuickChange II Site-Directed Mutagenesis Kit (Agilent Technologies), following the manufacturer’s instructions. The mutagenesis was performed on the plasmids encoding the Sph1 BiFC system ^37, 38^ and confirmed by a DNA sequencing program.

### Cell culture

#### Immortalized cell lines

Human Embryonic Kidney 293 (HEK) cells were cultured in Dulbecco’s Modified Eagle Medium (DMEM, Life Technologies-Invitrogen, USA), supplemented with 10% Fetal Bovine Serum Gold (FBS) (PAA) and 1% Penicillin-Streptomycin (PAN), and the cells were maintained at 37°C in an atmosphere of 5% CO2.

#### Primary cortical neurons

Primary midbrain neuronal cultures were prepared from E18 Wistar rat embryos (mixed sex) as described previously ^63^. The cells were seeded on poly-L-ornithine coverslips (Sigma-Aldrich) and cultured in a serum-free medium (Dulbecco’s Modified Eagle Medium/NutrientMixture F-12 [DMEM/F-12] consisting of N1 supplement, bovine serum albumin, glutamine (Sigma), and penicillin/streptomycin/mixture (Gibco). Every 3 days, one-third of the medium was removed and replaced with fresh media.

#### Primary hippocampal neurons

Primary mouse neurons were prepared from the hippocampus of embryonic day E16–E18 CD1 mouse embryos as previously described^35, 64^. All procedures were performed according to the National Institutes of Health Guide for the Care and Use of Experimental Animals and were approved by the University of Pennsylvania Institutional Animal Care and Use Committee. Dissociated hippocampal neurons were plated at 100,000 cells/well (24-well plate) in Neurobasal medium (Thermo Fisher) supplemented with B27 (Thermo Fisher), 2 mM GlutaMax (ThermoFisher), and 100 U/ml penicillin/streptomycin (ThermoFisher).

### Cell transfection

#### HEK cells

The day before transfection, 120 000 cells were plated in 12-well plates or in microscope glass bottom plates (iBidi) dishes, or 240 000 cells in 6-well plates (Corning). The cells were transfected with equimolar amounts of the plasmids (or otherwise indicated in the figure) using Rotifect Plus (Carl Roth) or Metafectene (Biotex), according to the manufacturer’s instructions. After 24 or 48 hours the cells were collected or stained for further analysis. For the competing assay, aSyn BiFC was first expressed, and after 24 hours, either empty vector, VN-Sph1 or Sph1-VC was expressed.

#### Primary cortical neurons

Neurons were transfected with Lipofectamine300 (ThermoFisher) at DIV 4 with equimolar amounts of the plasmids, following the manufacturer’s recommendation. The cells were cultured until DIV 6, after which the cells were fixed and stained.

#### Primary hippocampal neurons

Hippocampal neurons were transfected with calcium-phosphate method at DIV 4 as previously described ^65^. Briefly, 2× BES-buffered saline solution containing phosphate ions (50 mM BES, 280 mM NaCl, 1.5 mM Na_2_HPO_4_xH_2_O, pH 7.02) were mixed with calcium chloride solution (2.5 M) containing equimolar amounts of the plasmids. The cell growth media was collected and reserved, and OPTIMEM without adds where added to the cells. Cells were incubated with plasmid-calcium-phosphate coprecipitates for 40 minutes followed by media change (growth media previous collected). The cells were fixed and stained at DIV 6.

### Immunocytochemistry

After transfection, HEK cells, primary cortical or hippocampal neurons were fixed with 4% paraformaldehyde at room temperature (RT) for 10 minutes. After washing 3 times with 1xPBS, the cells were permeabilized for 20 minutes with 0.1% Triton X-100 (Sigma-Aldrich)/1xPBS. Then, the cells were blocked in 1.5% normal goat serum (PAA)/1xPBS for 1 hour and incubated with primary antibody. The primary antibodies used in this study were: anti-aSyn (610787, BD Biosciences), anti-Sph1 (sc-365741, Santa Cruz), anti-Giantin (ab80864, Abcam), anti-MAP2 (17490-1-AP, Proteintech) overnight (ON), and secondary antibody (Alexa Fluor 568 donkey anti-mouse IgG or Alexa Fluor 568 goat anti-rabbit IgG, (Life Technologies-Thermo Fisher Scientific) incubated for 2 hours at RT. Cells were stained with Hoechst 33258 (Life Technologies-Thermo Fisher Scientific) (1:5000 in DPBS) for 5 minutes, and the coverslips mounted for imaging.

For the organelle studies, HEK 293T were fixed and blocked as previously described, and the following primary antibodies were used: anti-Giantin (ab37266, Abcam), anti-PMP70 (SAB4200181, Sigma), AP-1 (610385, BD Transduction Laboratories), anti-TOM20 (sc-11415, Santa Cruz Biotech), anti-G3BP1 (WH0010146M1, Sigma), anti-DCP1B (13233, Cell Signaling Technology), anti-CKAP4 (AB_1731083), anti-LAMP1 (sc20011, Santa Cruz Biotechnology), anti-GAPDH (sc-47724, Santa Cruz Biotechnology). For immunofluorescence, we used the following secondary antibodies: anti-rabbit IgG Cy3-conjugated (Sigma-Aldrich C2306), anti-mouse IgG Cy3-conjugated (Sigma-Aldrich C2181), anti-rabbit IgG Cy5 conjugated (Invitrogen A10523), Anti-Mouse IgG H&L (Alexa Fluor 488) (Abcam).

### Quantification of Sph1-aSyn inclusions

At least fifty positively transfected cells *per* condition were quantified based on the presence or absence of inclusions. The results were expressed as the percentage of the total number of transfected cells obtained from three independent experiments for each mutation.

For the quantifications of the ratios, at least 300 cells *per* condition were used.

### FRAP analyses

The HEK cells were plated in iBidi dishes and transfected as mentioned previously. FRAP experiments were performed on a Leica 6000B microscope equipped with an incubator to maintain the 5% CO_2_ and 37°C, and with UGA-42 Firefly scanner-based systems (Rapp OptoElectronic, Germany).

The samples were initially imaged for three consecutive time-frames (pre-bleach-I_0_), and then a circular region of interest (ROI) was bleached with three iterations scan (30% of 405 nm laser power) for 227ms. A maximum of three inclusions *per* cell was bleached/recorded, and fluorescence recovery was measured every 20 seconds for 5 minutes.

To quantify inclusion intensity recovery over time, we used a customized script for Fiji (ImageJ) software (Wayne Rasband, NIH, Bethesda, MD, USA). Fluorescence intensities of the defined ROIs were measured, and the average intensities pre-bleach and after bleach were corrected according to the background. The average fluorescence in the bleached area I(t) at each time point t was used to normalize FRAP recovery curves.

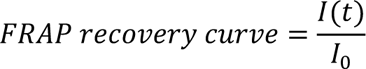

All nonlinear curve fitting and the statistical comparisons were performed using GraphPad Prism 4.0.

### Live cell imaging microscopy

For live cell imaging 4-well, iBidi or Cellview cell culture dish (Greiner Bio One) were used. Alternatively, cells were grown on glass slides (Marienfeld). Confocal images and movies were acquired using SP8 (Leica) confocal microscope equipped with a temperature and CO_2_ incubator, using a 60x PlanApo VC oil objective NA 1.40, and 406nm, 488nm, 561nm, and 640nm lasers. Image processing was performed using Fiji (ImageJ) software.

### Particle size analyses

The images were analyzed using a customized script in Fiji (ImageJ) software. Briefly, calibrated images (with pixels converted to micrometers) were used to automatically detect the particles. First, Otsu threshold was applied to convert the image to a binary image (black and white). Then, the minimum and maximum area sizes were set to exclude anything that is not the object of interest in the image. Once all the parameters were setup, the particle analysis was performed automatically. The data was blotted in a graph using BIN, which was performed in GraphPad Prism 4.0.

### Western blot analyses

Proteins were extracted using 1xPBS pH 7.4 with Protease Inhibitor Cocktail (1 tablet/10 mL) (Roche Diagnostics) and sonicated during 10 seconds at 10% power. Protein concentration was determined by Bradford assay (BioRad Laboratories), using an Infinite M200 PRO plate reader (Tecan, Lta). The sample were mixed with 5x Laemmli buffer (250 mM Tris pH 6.8, 10% SDS, 1.25% Bromophenol Blue, 5% β-Mercaptoethanol, 50% Glycerol), and denatured for 5 minutes at 95 °C. Precast gels were loaded with 30 μg protein, and the samples were separated on 4-20% or 4-15% Mini-PROTEAN TGX Precast Protein Gels (BioRad Laboratories). The gel was transferred to a PVDF membrane using the iBlot transfer system (ThermoFisher), according to the manufacturer’s instructions. Membranes were blocked with 3% (w/v) BSA (NZYTech), 1xTris Base Solution/0.05% Tween (TBST), followed by the primary antibodies diluted in 3% BSA/TBST ON at 4°C. The following antibodies were used: anti-aSyn (610787, BD Biosciences), anti-Sph1 (sc-365741, Santa Cruz), and anti-beta-actin (A2228, Sigma-Aldrich). After washing, the membranes were incubated for 2 hours with secondary antibody, anti-mouse IgG horseradish peroxidase labeled secondary antibody (GE Healthcare). Proteins were detected by ECL chemiluminescent detection system (Millipore) in Fusion FX (Vilber Lourmat).

### Native PAGE

48 hours after transfection, HEK cells were lysed in 1xPBS pH 7.4 containing Protease Inhibitor Cocktail tablet (1 tablet/10 mL) (Roche Diagnostics) and separated in 4–16% gradient Native pre-cast gel (SERVA Electrophoresis GmbH). Gels were run according to the manufacturer’s instructions, and transferred as previously described. The following antibodies used were: Syn1 for total aSyn (610787, BD Biosciences), MJFR-14-6-4-2-conformation specific (ab209538, Abcam) and 5G4-aggregated aSyn (MABN389, Merck).

### Proteinase K resistance

Cells were collected 48 hours after transfection in 1xPBS pH 7.4 with Protease and Phosphatase Inhibitor Cocktail (Roche Diagnostics). The lysates were sonicated for 10 seconds at 40–50% power, and then incubated with 2.5 μg/ml freshly prepared PK for different time periods (0, 5, 10 and 15 minutes). The digestion was stopped by adding the Laemmli buffer. The lysates were boiled at 95°C for 5 minutes, loaded onto 4-20% Mini-PROTEAN TGX Precast Protein Gels (BioRad Laboratories), and immunoblotted as mentioned above. The antibody used was Syn1 (610787, BD Biosciences).

### Cell viability

#### Cytotoxicity assay

Adenylate kinase release into cell media was measured 24 hours after transfection of HEK cells, using ToxiLight bioassay kit (Lonza) according to the manufacture’s recommendations.

#### Cell proliferation assay

The cell proliferation was evaluated after 24 hours of transfection using the AlamarBlue proliferation assay (BioRad Laboratories, USA). AlamarBlue was added at 1:10 ratio to the cells and quadruplicates of each condition was measured every hour on the infinite M200 fluorescence plate reader set to 37°C (Tecan Systems) with excitation 570 nm, emission 600 nm. The percentage reduction of AlamarBlue was calculated using the following formula:

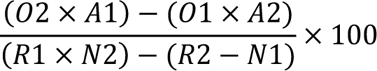

O1= Molar extinction coefficient of oxidized AlamarBlue at 570nm

O2= E of oxidized AlamarBlue at 600nm

R1= E of reduced AlamarBlue at 570nm

R2= E of reduced AlamarBlue at 600nm

N1= absorbance of negative control well (media with AlamarBlue but no cells) 570nm

N2= absorbance of negative control well (media with AlamarBlue but no cells) 600nm

#### Cell counting

48 hours after transfection, the HEK cells were trypsinized and cell viability was quantity using an automated cell counter (Countess from Invitrogen) at a 1:1 ratio of cells to trypan blue.

### Size exclusion chromatography and dot blot

Cells were harvested 48 hours after transfection in phosphate buffer (TBS with 0.5% Triton X-100), freshly supplemented with protease inhibitor cocktail, and centrifuged at 5 °C and 10,000g for 10 minutes. 1.5 mg of total protein, in a maximum volume of 500 µl, was filtered using a 0.45 µm spin-X centrifuge filter before loading onto a Superose 6 column (Superose 6 10/300GL, GE Healthcare, Sweden) and subsequent processed by SEC-HPLC (Äkta Purifier 10, GE Healthcare, Sweden). Each SEC-HPLC run was performed at a flow rate of 0.5 ml/min for 1.2 column volumes in ammonium acetate buffer (50 mM, pH 7.4). After the SEC-HPLC run, the collected fractions were boiled at 95°C for 5 minutes, centrifuged at 14,000g for 5 minutes, and loaded into a nitrocellulose membrane using a dot-blot apparatus. The immuno-signal was detected as described above, using the Syn1 antibody to detect total aSyn.

### Quantification of Golgi fragmentation

The Golgi morphology was assessed and scored into three groups as we previously published ^38, 40, 49^ (normal, diffused, and fragmented). Briefly, 48 hours after transfection, HEK cells were fixed and stained for Golgi apparatus as described in the previous section. For our analyses we discarded cells there were in metaphase (visible by the nuclear morphology). The morphology of the Golgi apparatus was assessed taking into consideration it shape and the area/distance to the nucleus: normal-compact/close to the nucleus; diffused: disperse/close to the nucleus; fragmented: dispersed throughout the cell. Three independent experiments were performed to determine the Golgi morphology of the cells expressing Sph1 and/or aSyn and the surrounding cells.

### Cell cycle arrest

For the double-thymidine cell cycle arrest, the cells were initially growth in medium containing 2 mM thymidine for 16 hours and then rinsed in 1xPBS, and maintained in growth medium for 8 hours. During this recovery period, the cells were transfected as before. Following this, the cells were exposed again to thymidine for an additional 16 hours before the final release of the cell cycle arrest. After 12 hours, the cells were rinsed, fixed, and prepared for analysis.

### Measurements of the area of the Golgi apparatus

To measure the Golgi area, we first converted our images (the same ones used for quantifying Golgi fragmentation) to 8 bits. Subsequently, we drew a box around the region of interest (the Golgi apparatus) and employed the Hull and Circle plugin from ImageJ to obtain classical shape descriptors, like circularity, convex area, convex perimeter, bounding box width, and height (as shown in supplementary Figure 2C). The convex area was chosen as the representative data, closely resembling the Golgi staining area distributed throughout the cytoplasm. These data were then plotted in a frequency distribution graph, in which we used relative frequencies to determine the percentage of values in each bin.

### Statistical analyses

The data were analyzed using GraphPad Prism 4 (San Diego California, USA) software. *P* values were calculated by two-tailed Student *t-test*.

For the analysis of organelles colocalization, three or more independent experiments were performed to obtain the data. *P* values were calculated by two-tailed Student *t-test,* or one-way ANOVA for samples following normal distribution determined by the Shapiro-Wilks test. The equality of variances was verified by Brown-Forsythe or F test. Mann-Whitney (2 groups), or Kruskal-Wallis (multiple groups) tests were used for samples that did not follow a normal distribution. The sample sizes were not predetermined.

Statistical significance was assessed where * corresponds to p < 0.05, ** corresponds to p < 0.01 and *** corresponds to p < 0.001.

## Data availability

All data are available within the Article and Supplementary Files, or available from the corresponding authors upon reasonable requests.

## Acknowledgements

This study was supported by Parkinson Fonds International, Germany. T.F.O. is supported by BMBF through the EU Joint Programme on Neurodegenerative Disease Research (JPND, www.jpnd.edu) project (OligoFit) and by the Deutsche Forschungsgemeinschaft (DFG, German Research Foundation) under Germany’s Excellence Strategy - EXC 2067/1-390729940, and by SFB1286 (B8). TA was supported by the HFSP Long-term Fellowship (LT000559/2021-L), EPFL Faculty, and the University of Southampton.

## Author Contributions

PIS assisted in the preparation of primary neurons. EG cloned Sph1 into the BiFC system and performed the HPLC and dot blot experiments. TA conducted the experiments shown in Figure 1C, 4D, Figure 5, Figure 1 A and C, supplementary data Figure 2C, supplementary data Figure 3, and supplementary data Figure 4A. DFL designed the study, performed the remaining experiments, and interpreted data. TFO designed the study and interpreted data. DFL and TFO wrote the manuscript. DM interpreted data. DM, DFL and TFO revised the manuscript and participated in discussions about the data.

## Supplementary figure legends

**Figure 1.**
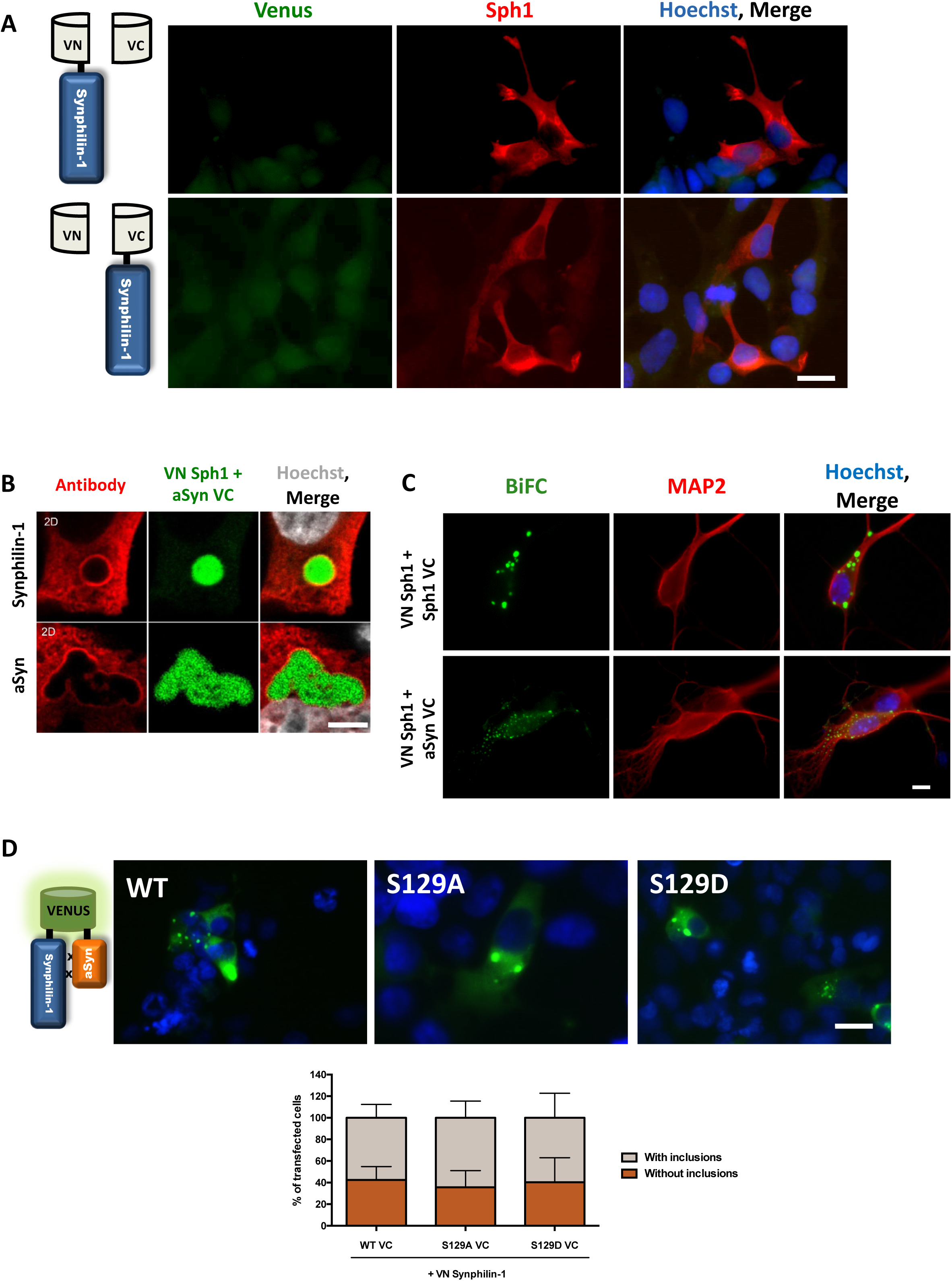
**A. BiFC controls.** No positive signal was observed for VN-Sph1 + -VC nor VN- + Sph1- VC. A diffuse and low signal was detectable for VN- + aSyn-VC. **B.** Confocal images showing a close detail of Sph1-aSyn inclusions (scale bar: 5 µm). **C. Primary hippocampal neurons exhibit Sph1-aSyn inclusions.** Transfected primary hippocampal neurons at DIV4, exhibit similar inclusions as HEK cells and primary cortical neurons at DIV6 (scale bar: 30 µm). **D. Phosphorylation does not change the inclusion pattern.** Blocking or promoting aSyn phosphorylation at S129 did not change the formation of Sph1-aSyn inclusions.

**Figure 2.**
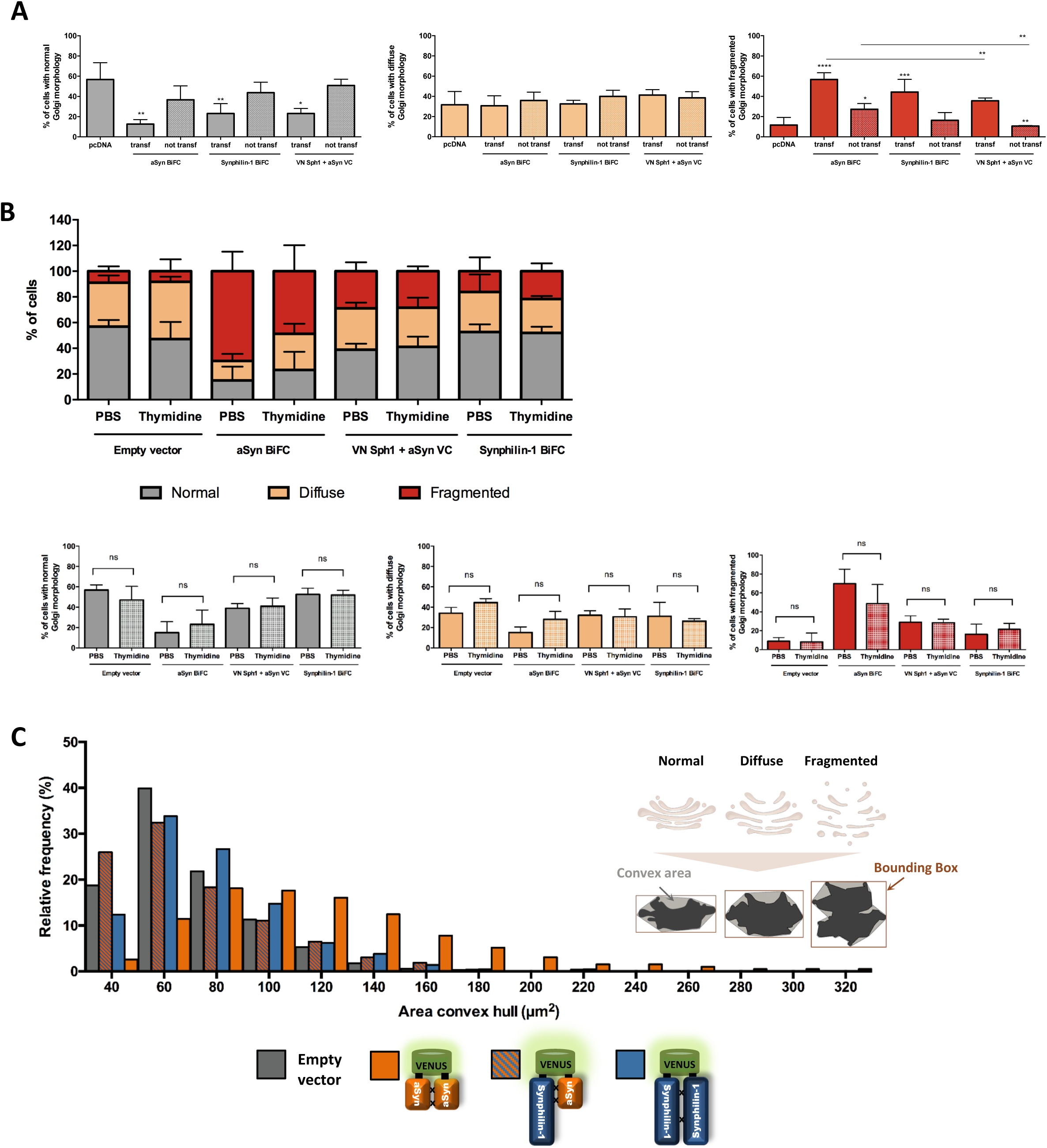
**A. Assessment of Golgi morphology. A.** Quantification of normal (gray), diffuse (yellow), and fragmented (red) Golgi showed that aSyn BiFC induced more fragmentation, but this effect can be contra balance with Sph1 expression. **B. Cell cycle arrest.** Thymidine-induced cell cycle arrest did not alter the overall toxic effects of aSyn on the Golgi apparatus. No observable differences were found between unsynchronized cells and those at the G1/S boundary. **C. Distribution of Golgi area**. aSyn BiFC occupies a larger area than VN-Sph1 + aSyn-VC and Sph1 BiFC. These results are in line with our observation/quantification. n=2.

**Figure 3.**
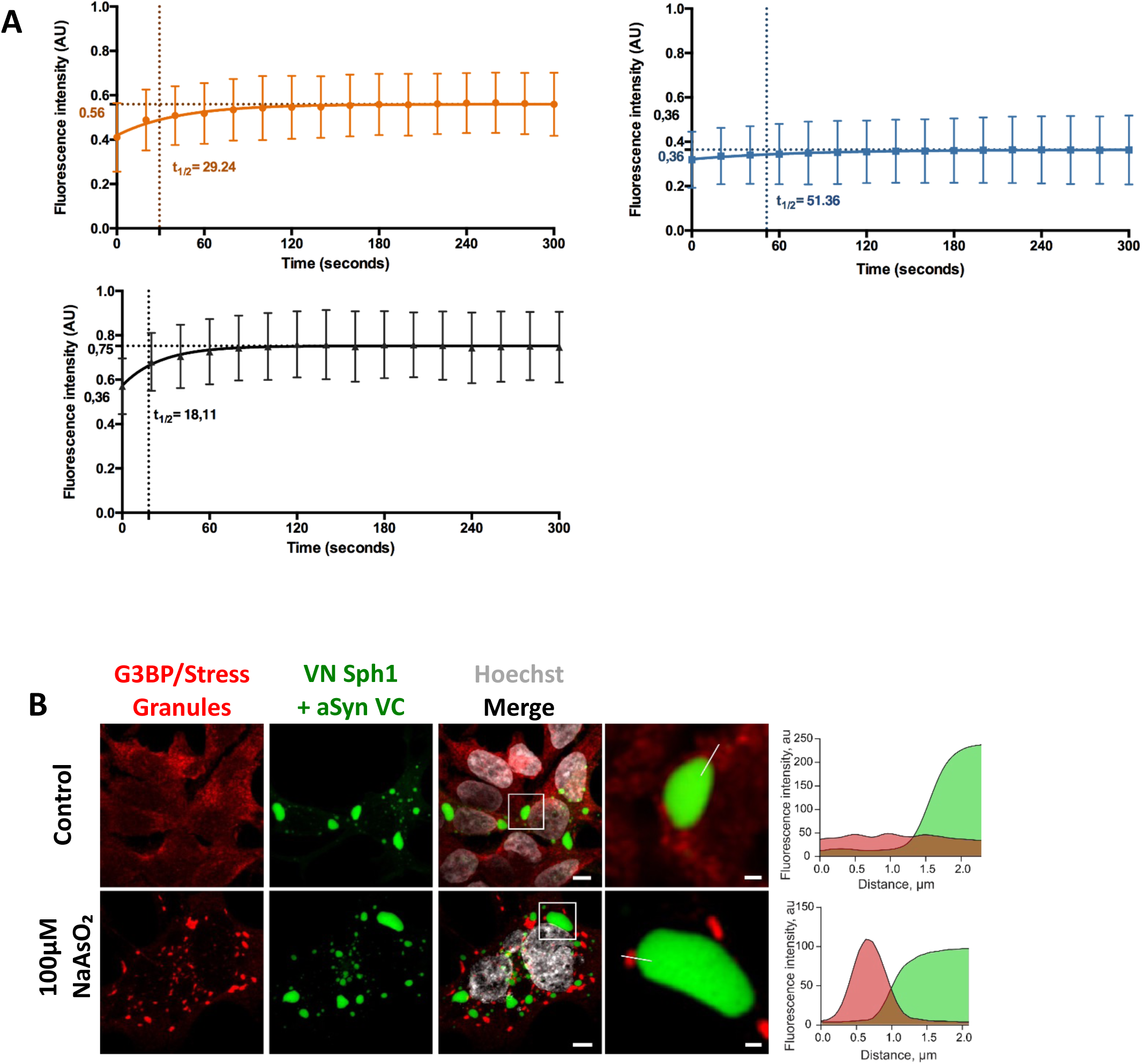
**A. Quantification of FRAP experiments.** The recovery of the fluorescence Venus signal was followed for 5 minutes, and the individual conditions were plotted in individual graphs showing the half-time value. **B. Stress granules do not localize with Sph1-aSyn.** Representative confocal images of cells expressing BiFC plasmids in control or arsenite (100µM 1 hour) conditions, stained with G3BP antibody. Fluorescence intensity profiles are shown (white line on the inset images). Scale bar of overall image: 5 µm and crop image: 1 µm.

**Figure 4.**
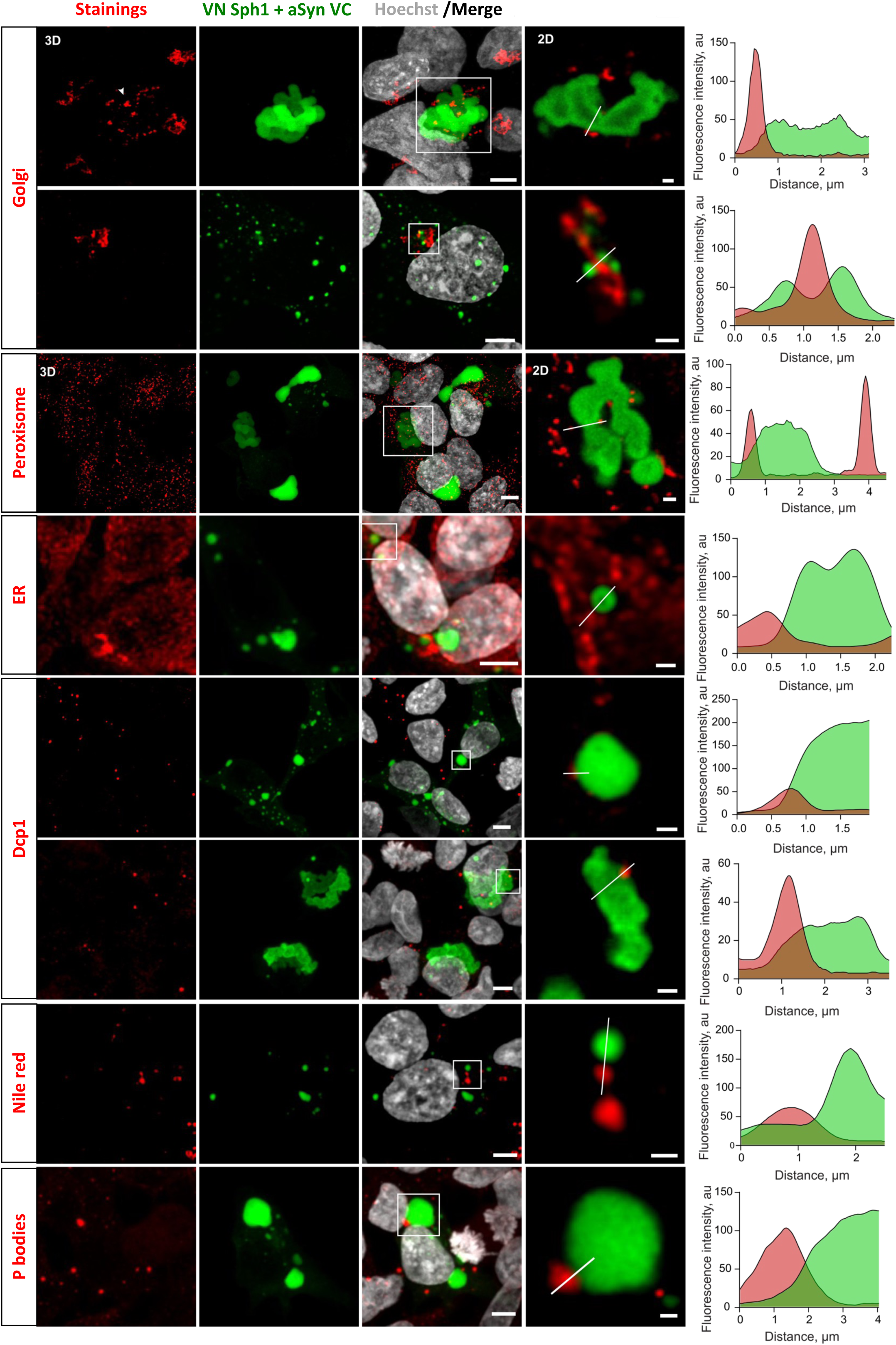
Organelle imaging. Several markers for membrane and non-membrane compartments were used to investigate the association of Sph1-aSyn inclusions with the organelle dynamics. Using confocal imaging, we found that inclusions did not co-localize with peroxisomes, ER, endoplasmic reticulum, lipid droplets, or P-bodies. Representative fluorescence intensity profiles through the inclusions (white line on the inset) are shown Scale bar of overall image: 5 µm and crop image: 1 µm.

**Figure 5.**
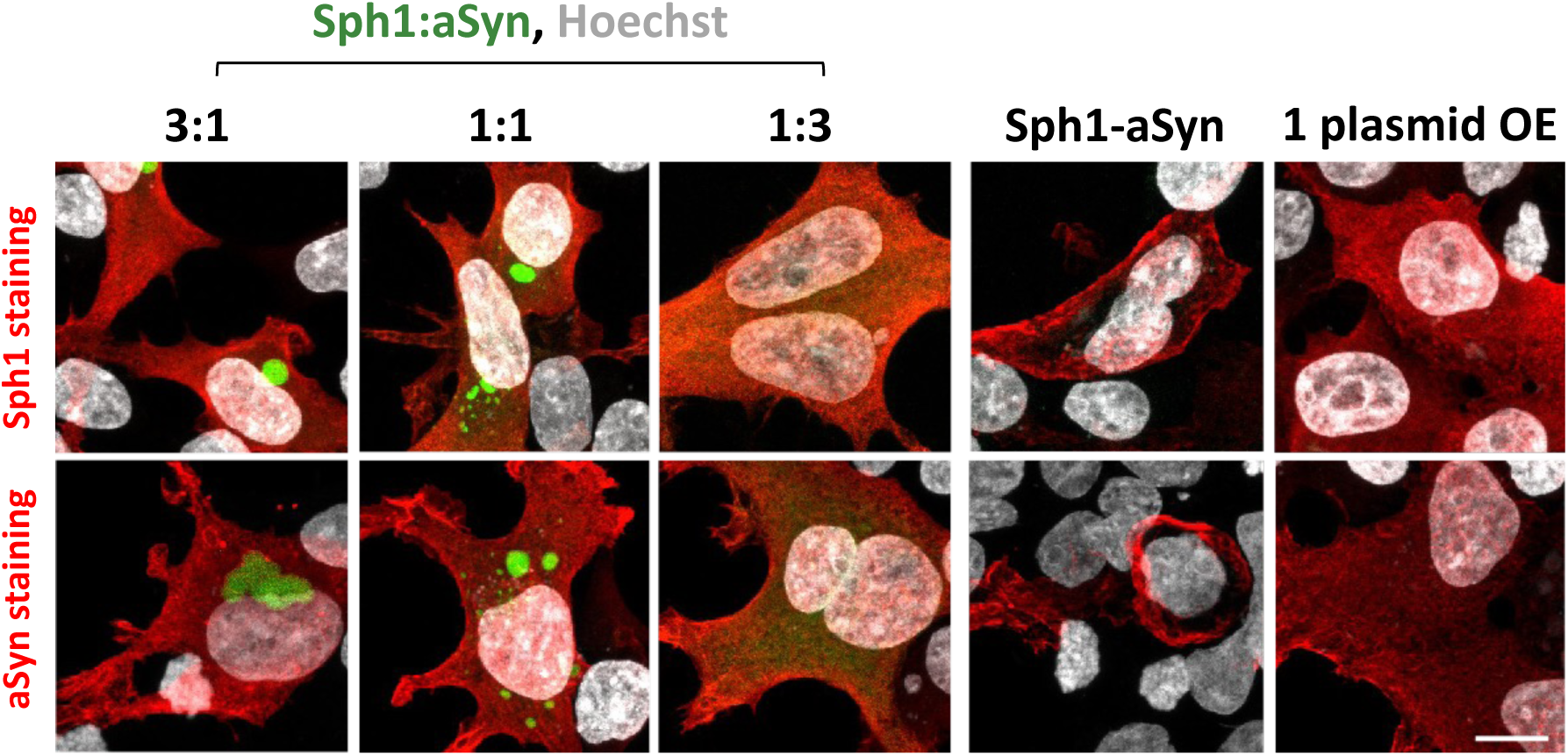
Sph1-aSyn localize with membranes. We constructed a plasmid in which we fused Sph1 and aSyn to guarantee similar expression levels for both proteins. We confirmed the presence of Sph1-aSyn at the membrane is independent of the expression ratio of Sph1 and aSyn, and their interaction is essential for their localization at the membrane. Scale bar: 5 µm.

